# Growth Condition Dependent Differences in Methylation Implies Transiently Differentiated DNA Methylation States in *E. coli*

**DOI:** 10.1101/2022.03.24.485589

**Authors:** Georgia L Breckell, Olin K Silander

## Abstract

DNA methylation in bacteria frequently serves as a simple immune system, allowing recognition of DNA from foreign sources, such as phages or selfish genetic elements. It is not well established whether methylation also frequently serves a more general epigenetic function, modifying bacterial phenotypes in a heritable manner. To address this question, here we use Oxford Nanopore sequencing to profile DNA modification marks in three natural isolates of *E. coli*. We first identify the DNA sequence motifs targeted by the methyltransferases in each strain. We then quantify the frequency of methylation at each of these motifs across the genome in different growth conditions. We find that motifs in specific regions of the genome consistently exhibit high or low levels of methylation. Furthermore, we show that there are replicable and consistent differences in methylated regions across different growth conditions. This suggests that during growth, *E. coli* transiently differentiates into distinct methylation states that depend on the growth state, raising the possibility that measuring DNA methylation alone can be used to infer bacterial growth states without additional information such as transcriptome or proteome data. These results provide new insights into the dynamics of methylation during bacterial growth, and provide evidence of differentiated cell states, a transient analogue to what is observed in the differentiation of cell types in multicellular organisms.

## Introduction

Cellular phenotypes are determined not only by genetic and environmental factors, but also epigenetic factors (heritable changes to the phenotype which are not caused by changes to the DNA sequence). In bacteria, epigenetic inheritance of phenotypes is known to occur via a range of mechanisms, including transgenerational inheritance of transcription factors or membrane transport proteins (Lambert and Kussell 2014; Kaiser et al. 2018), protein aggregates (Govers et al. 2018), or by covalent modifications to DNA, such as methylation (Sánchez-Romero and Casadesús 2020; Hale, van der Woude, and Low 1994). There are three types of covalent DNA modifications commonly found in bacteria: C^5^-methyl-cytosine (5mC), C^6^-methyl-adenine (6mA) and N^4^-methyl-cytosine (4mC) (Sánchez-Romero, Cota, and Casadesús 2015; Blow et al. 2016; Oliveira 2021; John Beaulaurier, Schadt, and Fang 2019). Methylation at these sites occurs via the action of DNA methyltransferases (Heard and Martienssen 2014; Jablonka and Raz 2009; Casadesús and Low 2006), which are ubiquitous across bacteria (Oliveira and Fang 2021).

Despite the ubiquity of DNA methylation, how often it serves an epigenetic function in bacteria is not well-established. In many cases, DNA methylation does not lead to different heritable phenotypes, and thus does not function as an epigenetic mark (Waldminghaus and Skarstad 2009; Skarstad, Boye, and Steen 1986; Collier 2009). However, a number of studies have established that DNA methylation can act to regulate cellular processes, including gene expression (D. Roberts et al. 1985; Seong, Han, and Sul 2021), sometimes in a heritable manner (Low, Weyand, and Mahan 2001; van der Woude, Hale, and Low 1998; Casadesús and Low 2006; Sánchez-Romero and Casadesús 2020). These modifications can have significant downstream phenotypic effects (Sánchez-Romero and Casadesús 2020; Park et al. 2019). Notably, in almost all well-established cases, when DNA methylation functions in an epigenetic manner, it is highly localised (e.g. at the operon-level) (Hale, van der Woude, and Low 1994), or even for a single site (Birkholz et al. 2022). One exception to this is a recent study, which suggested that genome-wide DNA methylation patterns differ between free-living and terminally differentiated bacteroids of the soil bacterium *Rhizobium leguminosarum* (Afonin et al. 2021).

To further probe possible epigenetic functions of DNA methylation in bacteria, here we characterise methylation patterns for three natural isolates of *E. coli* across a wide range of growth conditions. We profile DNA methylation using Oxford Nanopore (ONT) sequencing (Simpson et al. 2017; Rand et al. 2017), and show that by comparing samples of native methylated genomic DNA to whole genome amplified DNA it is possible to identify the expected methyltransferase binding motifs. We then use a quantitative approach to show that across the genome, methylation levels vary in a predictable fashion, and that levels of methylation differ between growth conditions. These data suggest that *E. coli* cells undergo environment-dependent transient differentiation into different methylation states during growth. These changes are not a reflection of cell cycle states, but instead are heritable changes that are gradually lost after growth ends. These results raise the possibility that in bacteria, growth states can be inferred solely by quantifying DNA methylation patterns, and that these patterns correspond to transiently differentiated epigenetic cell states.

## Results

### Determination of Methylation Motifs

We first sought to determine which methyltransferases were present in each of three natural isolates of *E. coli*, denoted here as SC419, SC452, and SC469 (Ishii et al. 2006). We found the adenine methyltransferase *dam* (which recognizes GATC motifs) and the cytosine methyltransferase *dcm* (which recognizes CCWGG motifs) in all three strains. We also found one of the adenine methyltransferases EcoKII or EcoGVI in each of the three strains. Both of these target the same motif, ATGCAT, and are present in most *E. coli* strains (Fang et al. 2012; Adzitey et al. 2020). We identified the methyltransferase EcoGIX in strains SC419 and SC469. EcoGIX is an adenine methyltransferase, with a loosely defined motif sequence (Fang et al. 2012; Forde et al. 2015). Finally, we identified EcoGVII in strain SC469, which is a close homologue of DAM (Fang et al. 2012), and recognises the same target motif.

To determine whether each of these methyltransferases was active we used ONT sequencing to identify genomic sites where DNA was modified. We sequenced native DNA which may contain modified bases, and whole genome amplified (WGA) DNA which contains few, if any, modifications. We generated at least 50-fold genomic coverage of ONT data from native DNA and at least 100-fold genomic coverage of ONT data from WGA DNA. (**Methods**). Note that these fold-coverage values are mean coverage values over the whole genome. To determine which genomic sites were modified we used a simple statistical approach implemented by Nanodisco (Tourancheau et al. 2021). Nanodisco uses the differences in the raw nanopore signals from each sample to assign a p-value to every position in the genome using a Mann-Whitney U-test.

We selected flanking regions from the 5,000 bases with the lowest p-values for input into MEME (Bailey et al. 2009) to identify motifs associated with modified bases. However, we found that in almost all cases, MEME identified only the cytosine methyltransferase DCM motif (CCWGG). We hypothesised that this was because methylated DCM motifs generally have smaller p-values than other motifs, due to larger signal deviations from unmethylated motifs. Because there are more than 13,000 DCM sites in each genome, the vast majority of the regions with low p-values would have been DCM sites, even when considering a very large number of sites (e.g., more than 10,000). We found that using a larger number of regions for input into MEME was computationally prohibitive. We thus randomly subsampled 100,000 base pairs (and associated p-values) from the genome (representing approximately 2% of the genome). From this subsample, we selected the flanking regions for the 5,000 base pairs with the lowest p-values for input into MEME.

For all three strains, MEME identified GATC and CCWGG as significant motifs (**Table 1**). These are the canonical motifs for the DAM and DCM methyltransferases, respectively, and we had bioinformatically identified both in all three strains. As these match the DAM and DCM motifs, we assumed that they contain C^6^-methyl-adenine (6mA) at the A position and C^5^-methyl-cytosine (5mC) at the second cytosine, respectively. Although we computationally identified the adenine methyltransferases EcoKII and EcoGVI in the three strains, we did not identify their target motif ATGCAT in any strains. We speculate that this is because methylated adenines are more difficult to identify (see above), and because this six-base pair motif is considerably rarer than the four-base pair motifs recognised by DAM and DCM. We also identified methyltransferase activity at two additional motifs, CCGG and GAGCC, in SC419 and SC452, respectively. Although there are no experimentally validated methyltransferases in the REBASE Gold database that are known to target these motifs, there are several putative type III R-M system methyltransferases that are thought to target these motifs. We mapped the sequences of each of these putative methyltransferases against each genome and identified a single genomic region in SC452 that matched all the putative GAGCC modifying methyltransferases (**Table 2**). This methyltransferase has a non-palindromic motif, and thus methylates only a single strand (Meisel et al. 1992). Surprisingly, we did not identify any CCGG-targeting methyltransferase in the SC419 genome. Finally, for the last two computationally identified methyltransferases, EcoGIX and EcoGVII, we could not unambiguously confirm any activity. This is not unexpected, as the EcoGIX motif is indefinite and the EcoGVII motif overlaps with DAM.

**Table 1.**
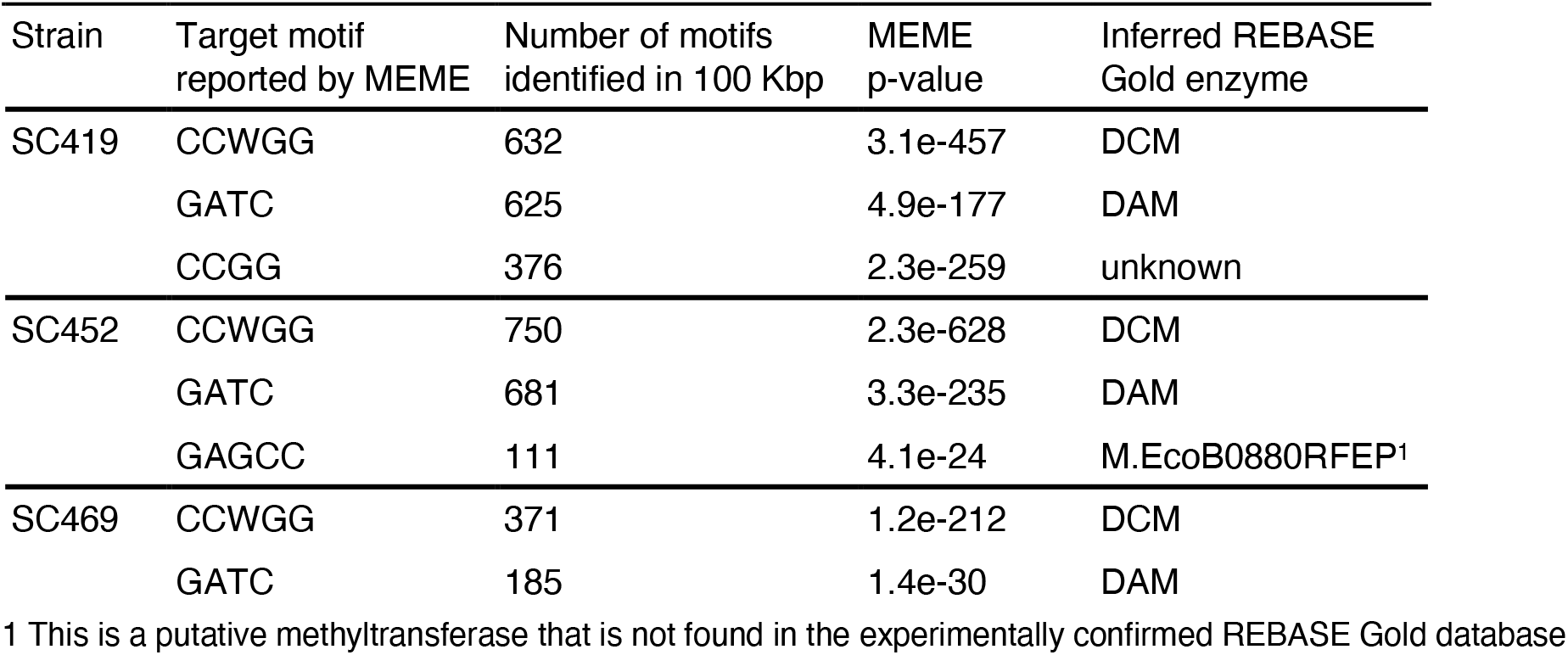
Matches between sequence motifs identified by MEME and REBASE Gold methyltransferases. Each row indicates the top three motifs as reported by MEME.

### Quantitative Analysis of Methylation Levels

We next sought to determine whether there was variation in the levels of methylation across the genome, or whether all regions were equally methylated. We focused only on the most commonly methylated motifs in each genome, GATC (containing methylated adenines via DAM) and CCWGG (containing methylated cytosines via DCM). Critically, the likelihood that a site is identified as methylated depends on the coverage of that site (**Fig. S1**). Thus, to increase the likelihood that all sites across the genome had an equal probability of being identified as methylated, we subsampled each of the ONT sequencing datasets to standardise coverage across the genome (**Methods**).

We then used Nanodisco to compare the native and WGA datasets for all three genomes, and for each known DAM and DCM motif site identified the lowest p-value from within the 3bp surrounding each motif (see **Methods**, *Quantification of methylation at individual sites*). These p-values should be indicative of the methylation status of a site, as they result from a Mann-Whitney U-test comparing the signal levels of modified and unmodified DNA. In addition, we hypothesised that sites at which all DNA molecules have a methylated nucleotide would have smaller p-values compared to sites at which only a small number of molecules are methylated, and that p-values are thus a quantitative indication of methylation status.

To directly test this hypothesis, we subsampled reads from the WGA data (which arises from fully unmethylated reads) to reach 50x coverage across the genome. We compared this WGA data with mixed native and WGA datasets having 50x coverage but consisting of 0%, 25%, 50%, 75% or 100% native reads. We expected that many of the native reads were fully methylated at DCM and DAM motifs. We then used Nanodisco to infer methylation status for all positions in the genome in these datasets with different ratios of WGA and native reads. We found a clear negative relationship between the fraction of native reads in the dataset and the associated p-values for each position (**Fig. 2**): as the fraction of native (possibly methylated) reads in the dataset increased, the p-values decreased. This indicates that the p-values returned by Nanodisco are correlated with the fraction of methylated molecules at a site and may provide quantitative insight into the fraction of molecules that are methylated at any DAM or DCM position in the genome. However, there are also clear complicating factors; for example, there is likely to be context-dependence of these p-values on the local nucleotide sequence.

**Fig. 1.**
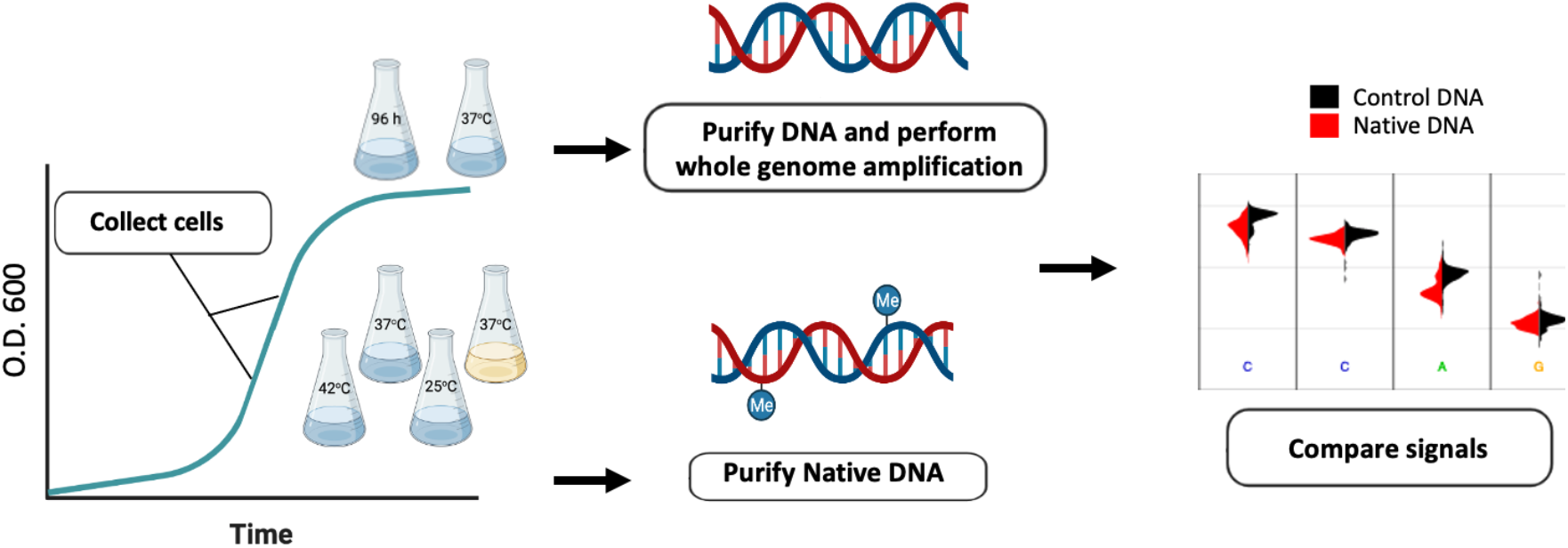
Experimental design for sampling native (possibly modified) and unmodified DNA. To sample native DNA, we grew cultures until exponential phase (for the minimal M9 media, rich LB media, 42ºC and 25ºC growth conditions); or late stationary phase (for the 96-hour growth condition). For whole genome amplification, we isolated DNA from early stationary phase (24 hours of growth). After purification of genomic DNA (and whole genome amplification when necessary), we sequenced the samples using the ONT platform. To infer DNA modifications, we compared the signals from native and WGA DNA using Nanodisco.

**Figure 2.**
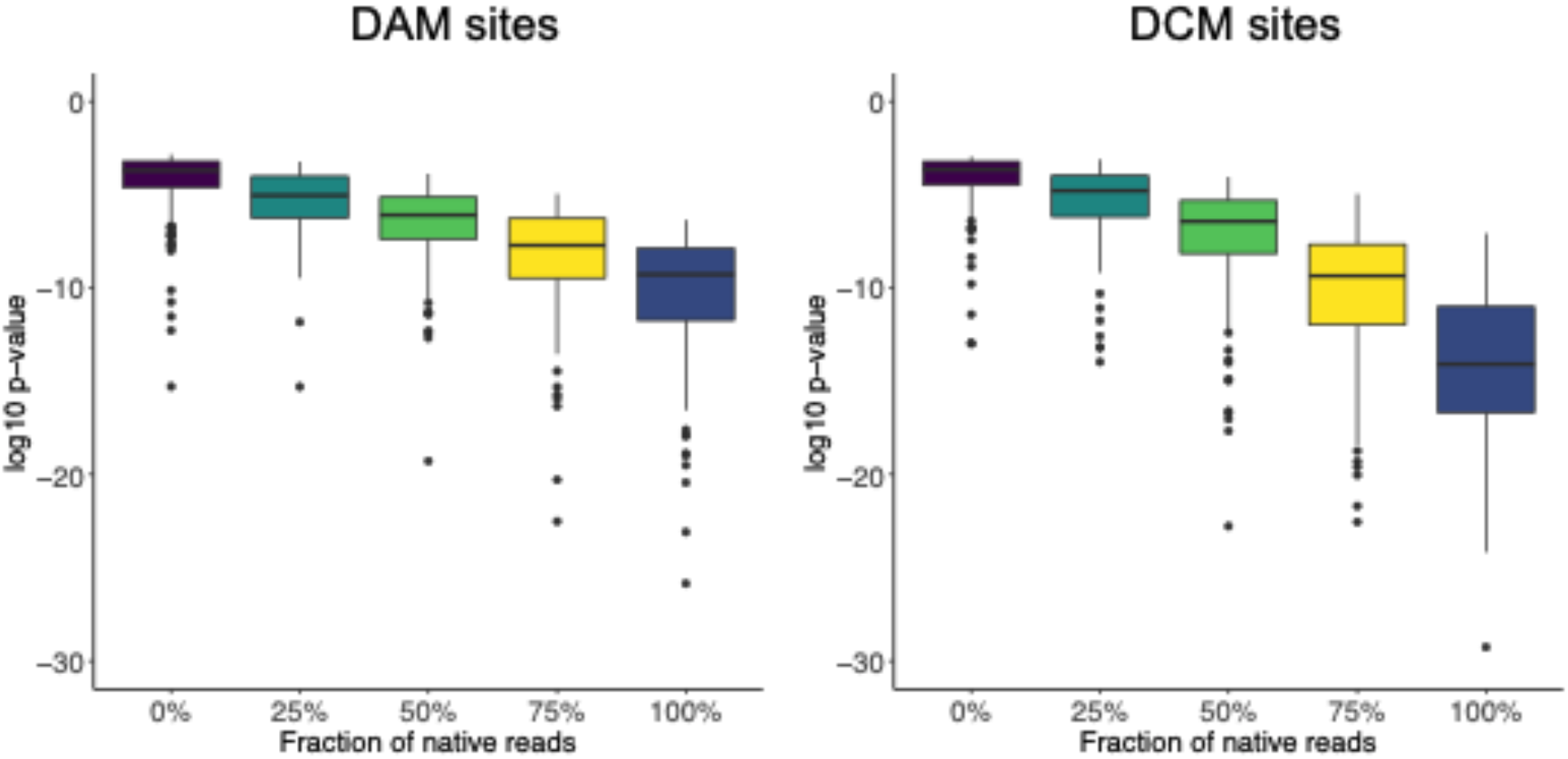
The p-values resulting from Mann-Whitney U-tests for signal deviations at DAM and DCM sites are correlated with the fraction of methylated molecules. We mixed known fractions of WGA reads (unmethylated) and native reads (possibly methylated) *in silico* and used Nanodisco to determine the p-value of a Mann Whitney U test at each position in the genome. We then determined the lowest p-value in a three bp window surrounding each hypothetically modified base in DAM (GATC) or DCM (GGCC) motif. For both methyltransferases, the sensitivity of the test increases as the fraction of native reads increases, with the DCM p-values decreasing to a much larger extent.

We then implemented a simple binary classification of DAM and DCM sites as being methylated or unmethylated (or less methylated) using a p-value cut-off (**Fig. S2** and **Fig. S3**). We placed this cut-off such that 10% of non-methylated sites were inferred as being methylated, analogous to implementing a false discovery rate of 0.1 (**Methods**; **Fig. S4** and **Fig. S5**). Although it would also be possible to implement a generative model specifying the fraction of molecules that are methylated at any one location in the genome, without a ground truth set of data for both unmethylated and methylated molecules, this is complicated. Thus, we use a simplistic binary classification. We note that, this division into methylated and unmethylated status for each site does not indicate definitively that a site is methylated or unmethylated. Rather, the division establishes that specific sites are more or less methylated (**Fig. 2**). We next used this classification of sites as methylated or unmethylated to test whether there were consistent differences in methylation rates across the genome or across growth conditions.

### Identification of Local and Global Methylation Patterns

To test for differences in methylation across growth conditions, for each strain we isolated DNA from cultures grown to exponential phase in five different conditions: two replicate cultures grown at 37ºC in minimal media (M9 glucose), one grown at 37ºC in LB broth (rich media), one grown at 25ºC in minimal media (low temperature stress), one grown at 42ºC in minimal media (heat stress), and one after 96 hours of growth in minimal media (late stationary phase). For each of these growth conditions, we performed the same p-value based analyses outlined above to determine whether DAM and DCM sites were classified as methylated or unmethylated.

We then used this data to look at large scale variation in methylation marks across the genome, based on both strain and growth environment. Rather than consider single sites, which exhibit considerable noise in being classified as methylated or unmethylated, we calculated the fraction of methylated sites in 10 Kbp windows across the genome (approximately 500 windows in total for a 5 Mbp genome; see **Methods**). Each of these windows contained approximately 40 DAM or DCM sites. We found that the fraction of sites classified as methylated within each 10 Kbp window varied by methyltransferase, strain, and environment (**Fig. 3**).

**Figure 3.**
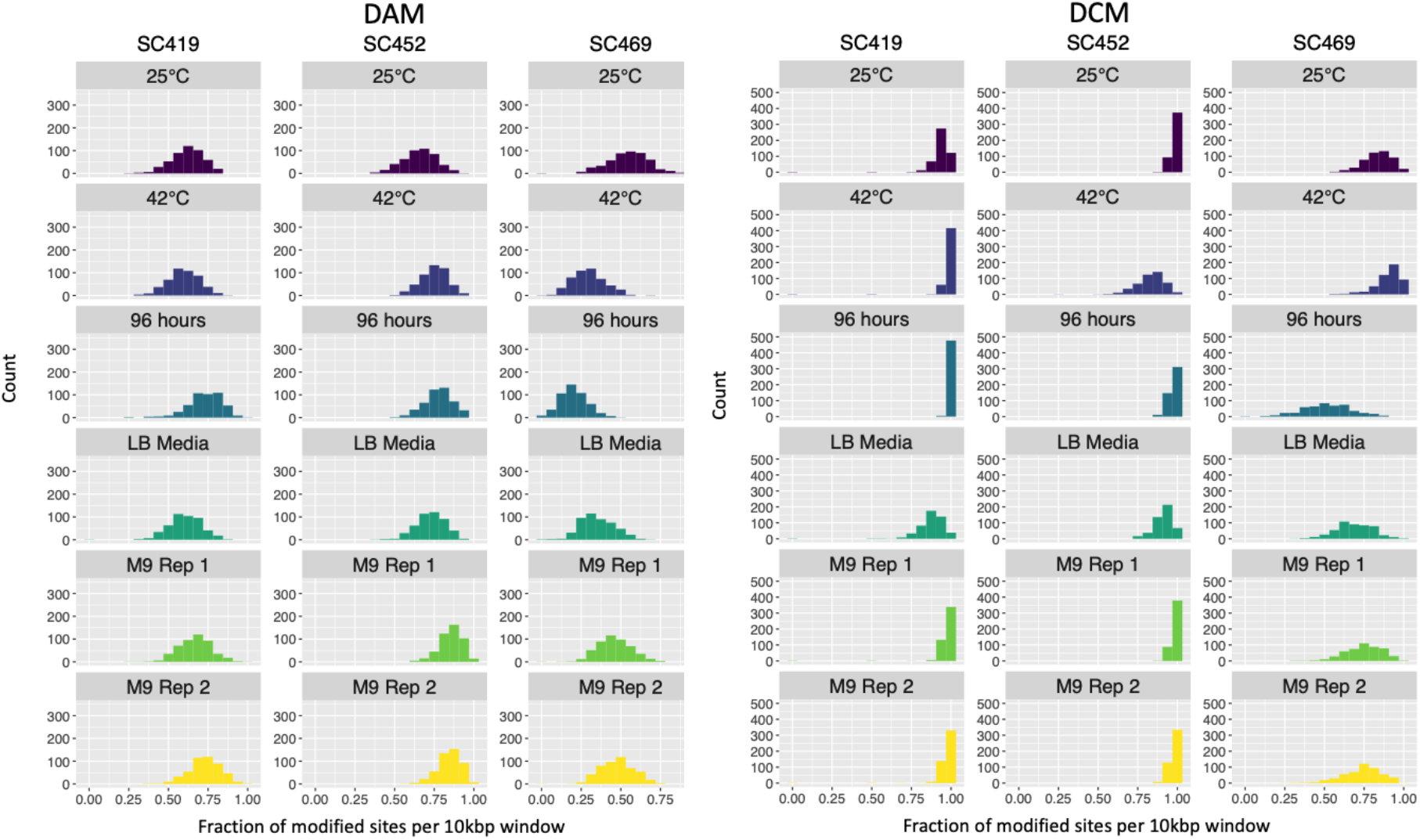
The fraction of DAM 6mA and DCM 5mC methylated sites within 10 Kbp windows varies according to strain and growth condition. The histograms in each panel indicate the distribution of 10 Kbp windows in which a certain fraction of sites are DAM (left panel) or DCM (right panel) methylated. This fraction ranges from almost 100% of all sites in all windows (e.g., for SC419 DCM in the 42ºC growth condition) to less than 50% of all sites in most windows (e.g., for SC469 DAM in the 42ºC growth condition). Except for the LB rich media sample, all cultures were grown in M9 minimal glucose media.

Overall, we inferred that a much higher fraction of DCM sites were methylated compared to DAM sites (**Fig 3**.). Part of this difference is likely due to the fact that the signal differences between methylated and unmethylated cytosines at DCM sites are much larger than between methylated and unmethylated adenines at DAM sites (**Fig. 2**). In these cases, it does not reflect biological differences but differences in the sensitivity of each statistical test. Nonetheless, we observed that in some growth conditions, a strain exhibited similar levels of methylation at both DCM and DAM sites (e.g., SC452 at 42ºC) whereas another strain in the same condition could exhibit different levels of methylation (e.g., SC469 at 42ºC). This indicates that it is unlikely that the lower levels of DAM methylation are due solely to decreased sensitivity, but instead to differences in the activity of each methyltransferase.

We also observed general strain-specific differences in methylation, for example, generally lower levels of both DCM and DAM methylation for SC469. However, it is difficult to determine whether this reflects real differences in methyltransferase activity between strains, or whether it is an artefact of the data analysis: for all cases, we inferred methylation status from a single unmethylated WGA dataset for each strain, and this in itself may cause differences in inferred methylation levels.

We next considered whether there were more localised patterns of methylation across the genome. To do this, we tested for correlations in the fraction of methylated sites within the 10 Kbp windows between growth conditions. Across different sets of growth conditions, we found that some 10 Kbp windows consistently had the majority of sites methylated, while other windows had many fewer sites methylated (**Fig. 4A**). It is possible that some of this is due to differences in coverage, as the relationship between inferred methylation status and coverage was not totally mitigated by our subsampling scheme (**Methods**). To minimise this dependence, we calculated the partial correlations in methylated fractions for each 10 Kbp window accounting for genome coverage (see **Methods**).

**Figure 4.**
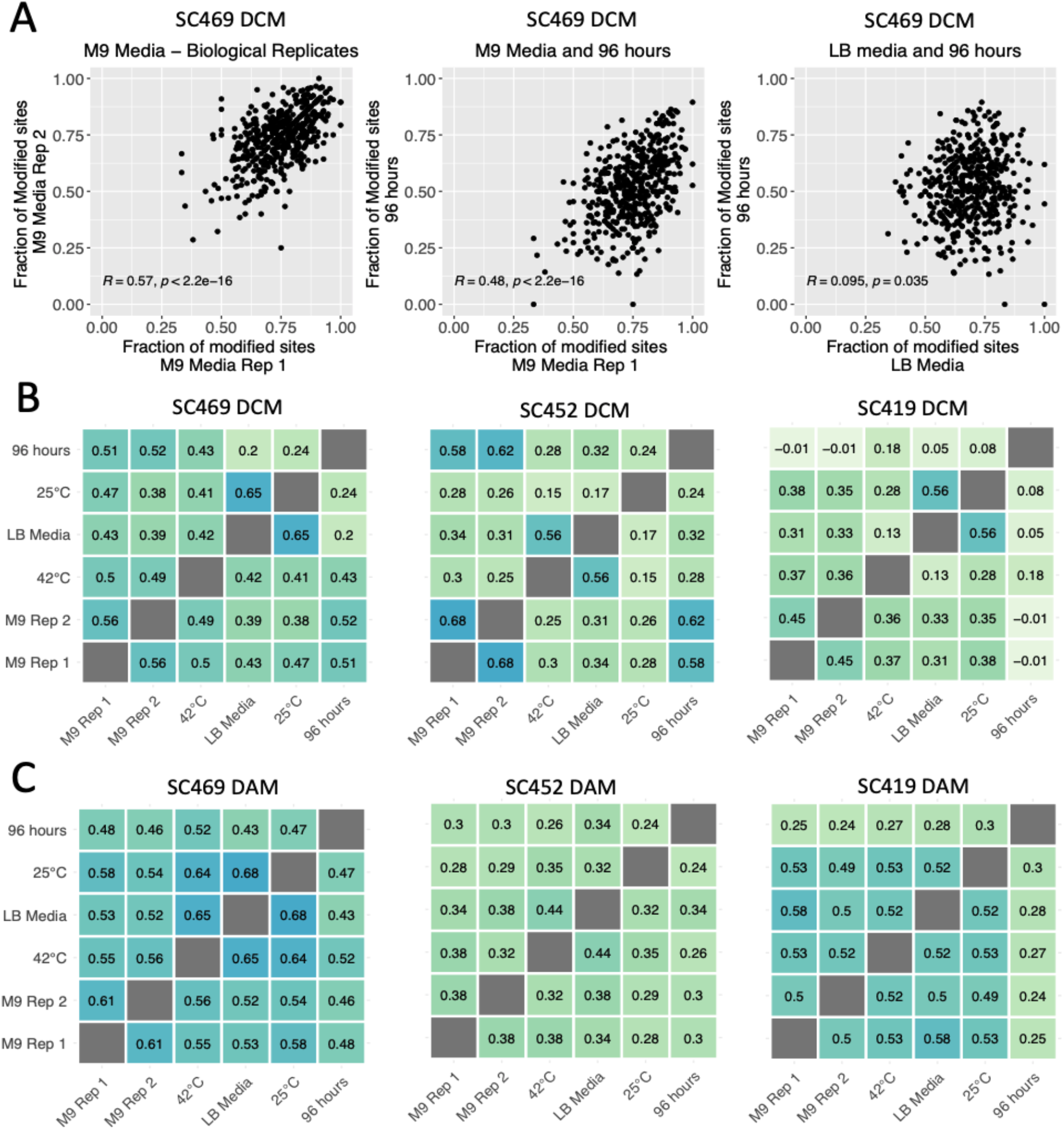
(A) The fraction of methylated sites in 10Kbp windows across the genome is correlated across growth conditions. The three panels indicate the fraction of methylated DCM sites within a 10 Kbp window that we inferred as methylated for strain SC469. We observed strong positive correlations in methylation patterns in replicate cultures of minimal M9 glucose media, slightly weaker correlations between M9 media and 96-hour stationary phase cultures, and almost no correlation between patterns in rich LB media and 96 hours stationary phase. Pearson partial correlations and corresponding p-values are indicated in each plot. **(B) Pairwise partial correlations in DAM and (C) DCM methylation patterns between all growth environments accounting for genome coverage**. Each panel shows all pairwise Pearson partial correlations between growth conditions in the fraction of methylated sites for all 10 Kbp windows in the genome, controlling for genome and WGA coverage in each of the growth conditions.

We calculated pairwise correlations in the fraction of methylated sites in 10 Kbp windows across the genome for both DAM and DCM in each strain across all pairs of growth conditions. We found replicable differences across the genome in methylation fractions (**Fig. 4**), with the correlations between some conditions being higher than others. Critically, we found that in all cases except one, the replicate cultures grown in M9 minimal glucose media at 37ºC exhibited the strongest correlation with the other M9 replicate. For example, for strain SC469 DCM the partial correlation between M9 replicates 1 and 2 was 0.56. The second strongest correlations for each were with cultures at 96 hours extended stationary phase (0.51 and 0.52 for replicates 1 and 2, respectively). Similarly, for SC469 DAM, the correlation between M9 replicates was 0.61. The second strongest correlations for each replicate were with growth at 25ºC (replicate 1, 0.58) and growth at 42ºC (replicate 2, 0.56).

This pattern, in which each M9 minimal media replicate correlated most strongly with the other replicate, extended to almost all strains and methyltransferases, with the single exception of DAM in strain SC419, for which methylation patterns correlated very similarly for all pairs of conditions (**Fig. 4C**, rightmost panel). As there are a total of six independent growth conditions, there is only a one in five chance that the two M9 replicates are most highly correlated. Thus, the likelihood that they would be the most highly correlated in almost all strains for both DCM and DAM strongly suggests there are growth-condition methylation states. Furthermore, these differences exist even when growth conditions differ only subtly (e.g., growth in minimal M9 glucose media at 37ºC versus M9 at 42ºC or growth in minimal media at 37ºC versus rich media at 37ºC).

In addition to high correlations between identical growth conditions, we often found consistent correlations in methylation status between different growth conditions. For example, the methylation patterns in the rich media LB condition (grown at 37ºC) often exhibited very strong correlations with methylation patterns in the minimal media 25ºC growth condition. In three cases (SC469 DCM, SC469 DAM, and SC419 DCM), these two conditions exhibited the strongest correlation of any pair of conditions. The convergent methylation states in these two conditions may be driven by similar changes in transcriptional activity, which could have an inhibitory effect on methylation.

The lowest levels of correlation we observed were for 96 hours extended stationary phase for strain SC419 DCM (**Fig. 4B**, rightmost panel). In some cases, the partial correlations were slightly negative. However, many of the 10 Kbp windows in this condition had almost 100% of all DAM sites methylated (**Fig 3**, right panel). Such low variability in methylation status means that strong correlations are difficult to obtain.

One explanation for the correlations in methylation fractions across growth conditions is that there are consistent long-range intragenomic correlations driven by periodicity in methylation, e.g. methylation fractions are generally lower at the origin of replication and higher at the terminus, or that there is transient methylation behind the replication fork (Anton and Roberts 2021). This would be apparent as long-range correlations in the fraction of methylated sites across the genome. For example, any two windows separated by a distance that is less than the periodicity should exhibit positive correlations. However, plotting the fraction of methylated sites across the genome revealed no strong long-range patterns (**Fig. 5**). To test for long-range patterns more systematically, we calculated correlations in the fraction of methylated sites within windows of increasing size, from 250 bp to 500 Kbp, separated by distances of increasing size, from 0 bp to 1 Mbp. This is similar to calculating an autocorrelation function, but for almost all step sizes (**Methods**). Again, we found no strong patterns of correlation between any windows larger than 5 Kbp, nor windows separated by more than 5 Kbp (**Fig. S7** and. **Fig. S8**). This suggests that short-range correlations dominate, and there are few long-range correlations in the fraction of methylated sites that are driven by factors such as higher levels of methylation at the terminus.

**Figure 5.**
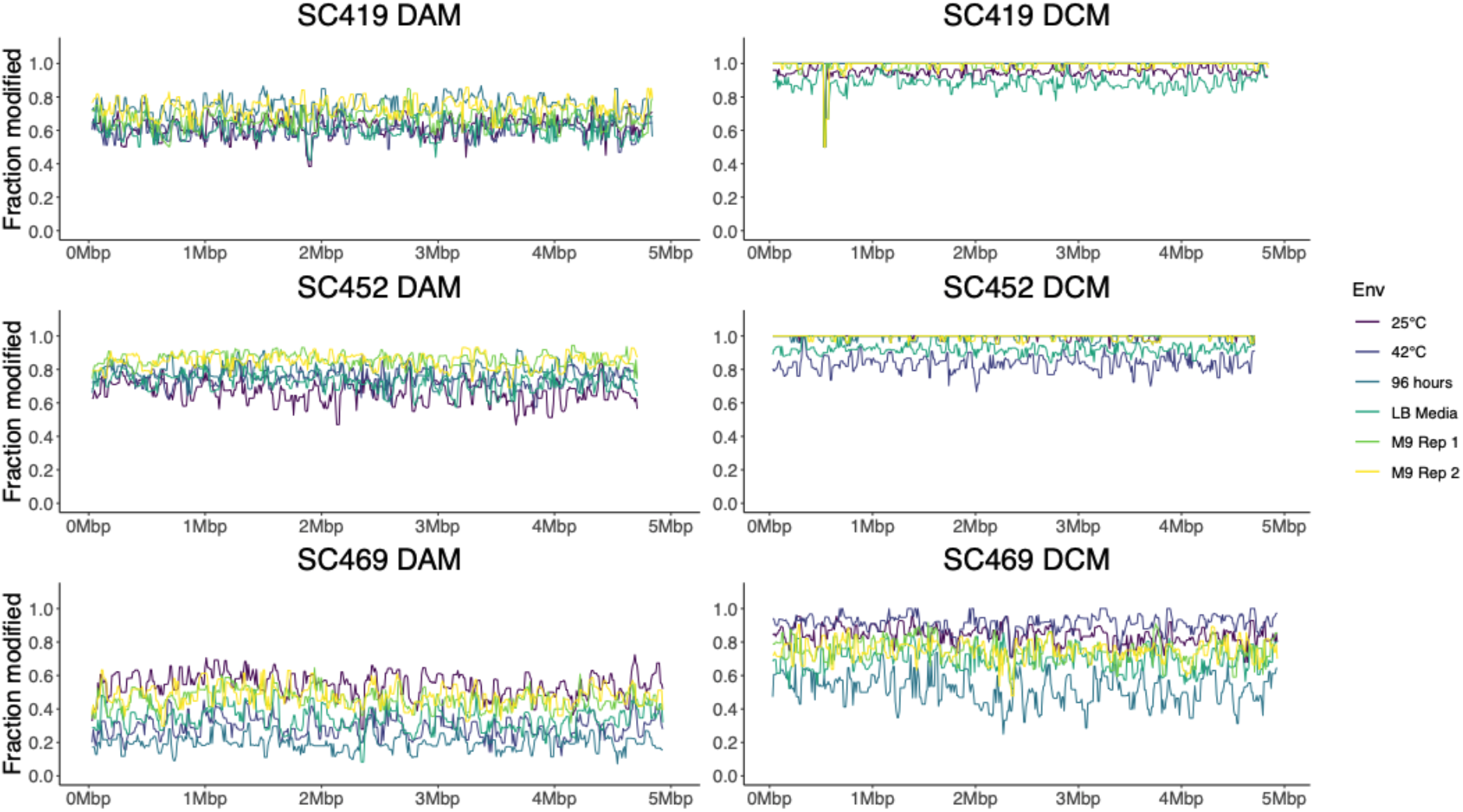
Genome wide patterns in the fraction of methylated sites. Each panel shows the fraction of methylated sites in 10 Kbp windows across the entire genome, with different growth conditions indicated in different colours. No long-range correlations, such as higher methylation at the replication terminus, were apparent.

## Discussion

Here we have identified DNA modifications in three *E. coli* natural isolates across a range of growth conditions using ONT sequencing. We have shown that it is possible to determine the motifs at which DNA modifications occur, and that these match the motifs expected given the restriction modification systems present in each genome. However, we also found one motif (CCGG) for which we could not identify a matching RM system; this motif may be modified by a novel methyltransferase.

Furthermore, we have shown that by using a simple binary classification of sites as methylated or unmethylated, it is possible to discern replicable and consistent differences in localised methylation frequency across the genome. The methylation patterns we have observed are dependent on growth conditions, with specific localised regions (on the order of thousands of kilobases) in the genome tending to be fully methylated, while others are less methylated. These conclusions differ from some previous work. A study on diverse strains of *M. tuberculosis* showed that most differences in methylation across the genome (as determined via SMRT sequencing) are due to stochasticity in intracellular methylation, rather than consistent differences between cells in methylation rates. Consistent differences between loci in methylation (hypomethylation) were found to be exceedingly rare, on the order of 10 to 20 sites across the genome (Modlin et al. 2020). Other work has also shown that methylation remains remarkably consistent across different growth conditions, including antibiotic stress (Cohen et al. 2016) and over the growth cycle (Payelleville et al. 2018). A significant difference between these latter two studies and the data we present here is the inclusion of methylation at DCM sites (CCWGG) in addition to DAM sites (DAM). Indeed, the most notable methylation patterns that we find – although subtle – are due to differences at DCM sites (**Fig. 4B**). Differential methylation at DCM sites has been connected to major changes in ribosomal gene regulation (Militello et al. 2012).

Critical to our proposal that these methylation patterns have epigenetic effects is that DNA methylation is heritable. Sites at which both the top and bottom strand are methylated will impart hemimethylated strands to both daughter cells, which will become fully methylated by “maintenance” methyltransferases (Anton and Roberts 2021); sites that are hemimethylated will impart one hemimethylated strand to one daughter cell and one unmethylated strand, which is more likely to remain unmethylated. This means that mother cells with methylation at a certain genomic location will have daughter cells that are also methylated at that location, but this will vary across daughter cells. Thus, if methylation affects phenotype, and methylation varies between individual cells in a population, then it acts as an epigenetic mark for the instances we have described here.

It is possible that there are unrecognised causes that drive some of the inferred differences in methylation status across the genome. For example, subtle differences in nucleotide context affect both the activity of the methyltransferase and the deviations in ONT signal. This undoubtedly influences our ability to accurately infer methylation status. However, we do not expect these differences to be dependent on growth conditions. Thus, the fact that we find both higher correlations between identical growth conditions, and consistently higher correlations between specific pairs of growth conditions (e.g., rich media (LB) at 37ºC and M9 minimal glucose media at 25ºC), suggest that nucleotide context is not the only force driving this correlation in methylation states. Additional work is required to test the repeatability of methylation patterns in different conditions, and whether other divergent growth conditions, for example antibiotic stresses or additional heat stress, lead to greater differences in methylation patterns. Similarly, methylation patterns should converge as growth conditions converge - for example we would expect more similar patterns comparing methylation during growth at 37ºC and 39ºC than to 42ºC. Again, more experimentation is needed here.

In eukaryotes, it is well-established that methylation affects gene expression (Song et al. 2005; Vanderkraats et al. 2013), and thus cell phenotypes. Here we have shown that methylation patterns are consistent and replicable in different growth conditions in *E. coli*. In addition, for identical growth conditions (in the data here, M9 minimal glucose media), there are strong correlations in which specific regions of the genome are methylated. There are two readily apparent explanations for these results. Either growth phenotypes affect patterns of methylation, or methylation patterns affect growth phenotypes (or both). We propose that it is likely that (as with eukaryotic cells) methylation affects gene expression in *E. coli* in different growth environments, although we have not established causation (Chen et al. 2018). This connection between methylation and transcriptional regulation has been proposed previously (Beaulaurier et al. 2015), and there are data that both support (Gaultney et al. 2020) and refute the connection (Mehershahi and Chen 2021). However, we note that there are many other well-established instances in which this causal direction has been established (Sánchez-Romero and Casadesús 2020).

Regardless of whether methylation functions as an epigenetic mark, and regardless of its causality, we have shown that just as bacterial cells undergo transient differentiation into different growth phenotypes, they also undergo transient differentiation into distinct methylation states. As we have not used synchronised cultures, it is unlikely that the correlated methylation is due to synchrony in the cell cycle that differs between growth conditions. This is further supported by the fact that we have shown that correlations do not arise because of short- or long-range correlation in methylation fractions (e.g., differences in methylation at the chromosomal replication ori or terminus). Rather, these correlations arise from localised differences across the chromosome.

This work raises the possibility of discerning bacterial growth states without measuring cell physiology or quantifying the transcriptome, similar to what can be done for differentiated eukaryotic cells. We propose that with sufficiently long reads and precise measurements, it will be possible to quantify methylation states across single molecules, and from there infer the growth state of a cell from which a particular DNA molecule has originated. In addition, with more nuanced model-based or machine learning analyses, it may be possible to assign genomic methylation patterns more specifically to specific growth states. This contrasts with more standard approaches such as single-cell transcriptome profiling, which is often of limited use in bacteria given the extremely small number of transcripts contained in most cells.

## Methods

### Bacterial Growth

We grew overnight cultures from single colonies for each natural isolate in 3mL of liquid LB media at 37°C. We then inoculated 75mL of the relevant growth media (either LB or M9 minimal media with 0.2% glucose) in a 250ml Erlenmeyer Flask with 75uL of overnight culture. We grew these at the relevant temperature (37°C, 25°C, 42°C) until an OD600 between 0.4 and 0.5 was reached, or for 24 hours or 96 hours (for WGA and late stationary phase samples). 5ml of media was removed into a 15ml falcon tube and the cells were pelleted by centrifugation at 14,000 RPM for four minutes. We removed the media and spun the cells for an additional two minutes, after which we pipetted off any remaining media. We stored the cell pellets at -20°C until DNA extraction.

### DNA extraction and whole genome amplification

We extracted DNA using the Promega Wizard DNA extraction kit following the gram-negative bacterial extraction protocol. We performed whole genome amplification (WGA) using the Qiagen RepliG kit according to the manufacturer’s protocol. We used a Qubit fluorometer to measure DNA concentration, ensuring that each sample had sufficient DNA for a ligation library prep without further concentrating the sample. We measured DNA purity with a Nanodrop. For all samples, the 260/230 and 280/230 ratios were between 1.5 and 2.3. We stored DNA at -20°C until library prep and sequencing.

### Library preparation and DNA sequencing

We prepared ONT sequencing libraries for both the WGA and native DNA using either the SQK-LSK109 kit with barcode expansion kit EXP-NBD104 or the SQK-RBK004 kit. For the SQK-LSK109 kit we followed the manufacturer’s protocol with no modifications. We modified the SQK-RBK004 protocol as follows: we eluted the samples off Agencourt Ampure XP beads using TE buffer pre-warmed to 50°C; we performed the elution itself at 50°C; and we increased the incubation time for elution to 10 minutes.

We performed ONT sequencing on a MinION Mk1B device using R9.4.1 flowcells. We used eight flowcells in total (two with SQK-RBK004 libraries and six with SQK-LSK109 libraries), with 12 samples run per flow cell. One additional flow cell was used to produce an additional 1 Gbp for a single sample that had low coverage. For each sequencing run, we demultiplexed and basecalled using Guppy v4.2.2.

For quantitative analysis of methylation, we subsampled all WGA and native sequencing reads to ensure even coverage across the genome using the following strategy: for each sample, we mapped all reads onto the relevant reference genome and determined the lowest 5th percentile of coverage over all samples, excluding the 96-hour sample, which had lower coverage for all strains (see below). For the 96-hour samples, we calculated the 5th percentile of coverage only for those samples, rather than across all samples.

We then standardised coverage across the chromosomal contig at this 5th percentile level. We first calculated the mean read length for each dataset. We then divided the genome into 10 Kbp windows and sampled an appropriate number of reads originating within each window such that the read length and the target coverage matched (e.g., if mean read length was 2 Kbp and the target coverage was 100X, then we selected 500 reads originating within the 10 Kbp window). We then mapped all reads back onto the genome to confirm that we had reached the coverage targets. If the target coverage was not achieved (for example due to irregularities in the read length distribution), the mean read length was adjusted to represent the mapped reads and reads were resampled. We then used the ONT-fast5-api to extract the corresponding fast5 reads for each dataset (see GitHub).

### Identification of methyltransferases

We previously produced reference-level genomes for each strain (Breckell and Silander 2020) using Prokka (Seemann 2014). We identified methyltransferases by using bwa mem (Li 2013) to map all restriction enzymes and methyltransferase enzymes in the REBASE Gold database (R. J. Roberts et al. 2010) to each strain. The REBASE Gold database contains only experimentally validated methyltransferase and restriction modification systems. We filtered the alignments to include only those genes which aligned for more than 97% of their length.

### DNA modification analyses

#### Detection of modified sites using Nanodisco

We used Nanodisco to detect DNA methylation (Tourancheau et al. 2021) with the recommended default settings. We processed fast5 reads from both WGA and native DNA samples separately with the Nanodisco pre-process command before running the Nanodisco difference command to calculate differences in the WGA and native DNA signals at each position. We used the Nanodisco merge command to create a single output file containing the native and WGA coverage for each genomic location, the mean signal difference and U- and t-test p-values reporting the significance of the signal difference at each site.

#### Quantification of methylation at individual sites

The Nanodisco output includes a p-value of a two-tailed Mann-Whitney U-test for each site indicating whether the signal at that site differs between the modified and unmodified samples. However, this p-value is not necessarily lowest at the actual point of modification, as the nanopore detects five bases at once, and the methylation can affect the signal in unpredictable ways. For example, many bases that were identified as having signals that differed between native and WGA DNA were not highest at the expected cytosine position within GATC motifs. To ensure we identified methylated motifs, we first identified all motif locations (DCM and DAM) in the genome (CCWGG and GATC, respectively), and then identified the lowest p-value out of the focal base and either neighbouring base. We used this p-value as an indication of whether a CCWGG or GATC site was methylated.

To account for false positive identification of modified sites, we used the p-values from above for the DCM and DAM sites located in the first 1 Mbp of the genome. We also identified an equal number of random locations in the first 1 Mbp of the genome, and identified the lowest p-value of each random bp or either neighbouring bp. We performed this analysis only in the first 1 Mbp of the genome to minimise computational effort; it is highly unlikely that this has any effect on the results. This resulted in a set of p-values for possibly methylated sites within each target motif, and likely unmethylated random sites. We used the p-values from the random sites to establish a null distribution of p-values for unmethylated bases. We designated all DAM and DCM sites with p-values lower than the 10th percentile of the null distribution as methylated (**Fig. S1**). All other DAM and DCM sites we designated as unmethylated. The precise implementation of this method is available through the GitHub repository indicated above.

#### Correlation in methylation fractions

To calculate correlations in the fraction of methylated sites, we first determined the number of DAM or DCM binding motifs within each 10 Kbp window for each genome. We used this as an estimate for the number of potential DAM or DCM modifications and then calculated the fraction of DAM or DCM sites which we experimentally identified as modified in each window. We calculated the correlation between the fraction of modified sites in each window as a Pearson correlation or a partial correlation accounting for sequencing coverage, as sequencing coverage affects the likelihood that a site will be detected as modified.

#### Genome wide methylation patterns

We assessed genome wide methylation patterns by comparing the fraction of known sites vs modified sites in windows across 10 Kbp windows in the genome. We discarded any regions that contained no DAM or DCM sites, as this would result in a division-by-zero problem. For the normalised data presented in **Fig. S6**, we simply divided the fraction of methylated sites in each window by the mean of all windows across the genome.

## Author statements

### Authors and Contributors

Conceptualisation: GLB and OKS. Methodology: GLB and OKS. Investigation: GLB. Writing – Original Draft Preparation: GLB and OKS. Writing – Review and Editing: GLB and OKS. Visualisation: GLB (lead) and OKS (supporting). Supervision: OKS. Funding: OKS.

## Acknowledgments

Thank you to N. Freed for assistance with whole genome amplification and Oxford Nanopore sequencing, B. Morampalli for fruitful discussions on inferring methylation, A. Sulit and M. Vklová for help with R and Python debugging, and A. Tourancheau for advice on using Nanodisco.

## Supplementary Figures

**Supplementary Figure S1.**
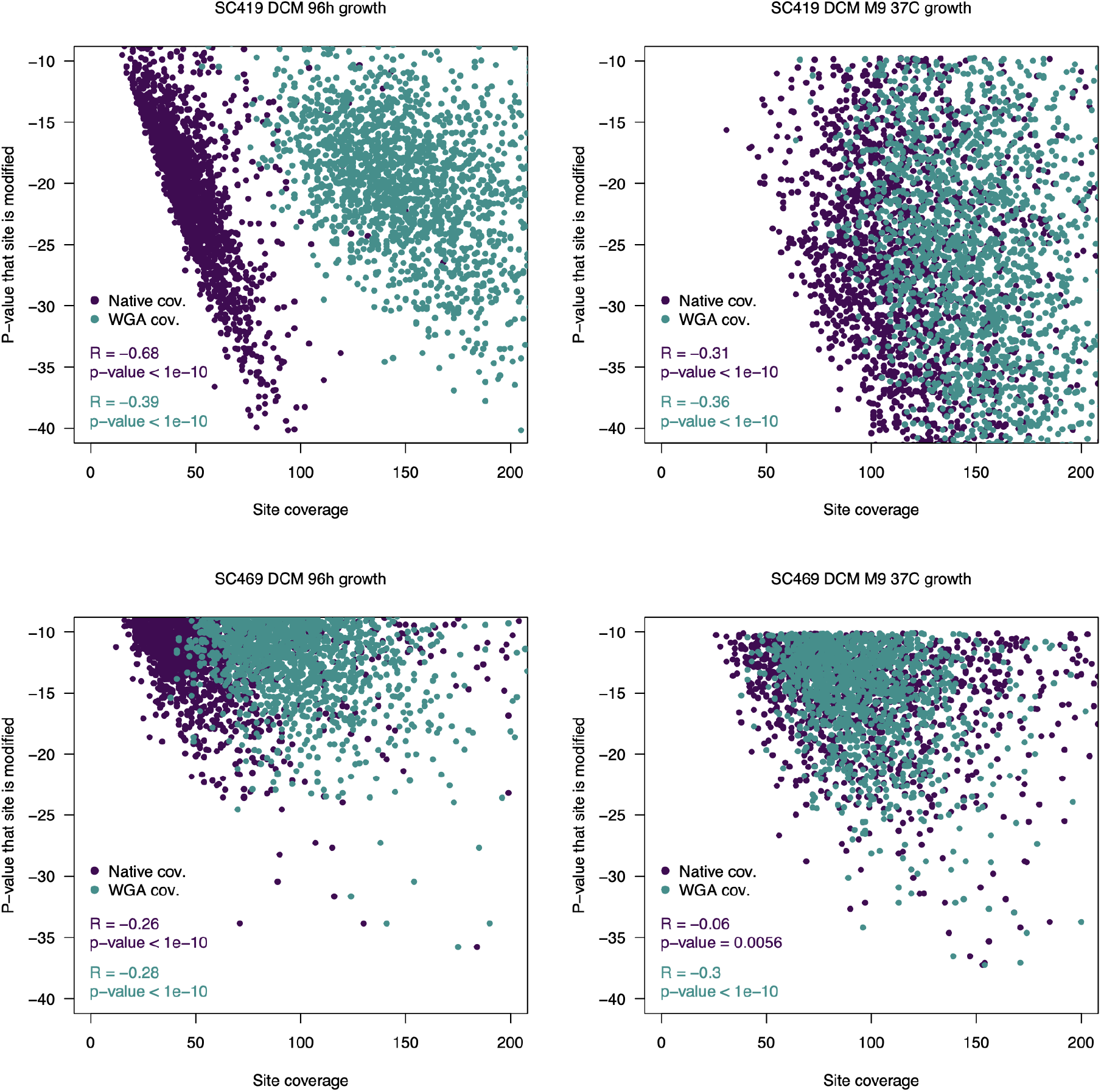
Correlation between coverage and the Nanodisco-derived p-values. Each point indicates the coverage at individual DAM or DCM sites and the p-value of the Nanodisco Mann-Whitney U-test. There is a clear relationship between the likelihood the p-value returned by Nanodisco (indicating a site is likely modified) and the coverage at that site, with both the coverage of the native DNA sample and the WGA sample affecting the test implemented by Nanodisco. The four examples above are all for DCM sites in two strains and two growth conditions for each. In all plots, only the sites that have p-values significantly lower than the null model background are shown. The native coverage at these sites is shown in purple; the WGA coverage at these same sites is in blue. For both native and WGA coverage, there is a strong negative correlation - sites with higher coverage have a lower p-value and a higher probability of being identified as methylated, although this differs between datasets. For example, there is only a weak relationship (R = -0.06) between native coverage and the p-value to the test in the SC469 DCM dataset.

**Supplementary Figure S2.**
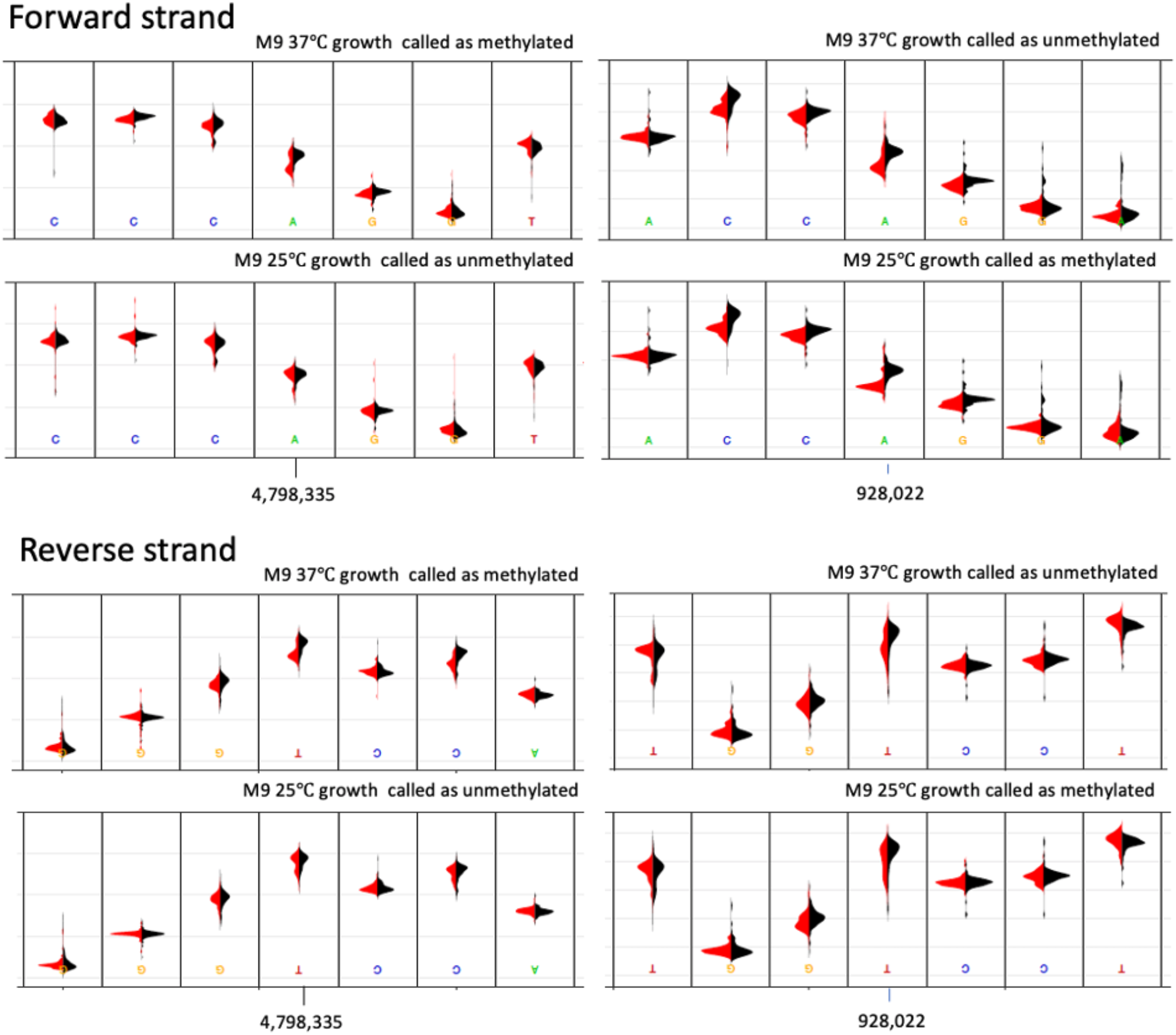
Raw nanopore signal distributions on the forward and reverse strands at identical genomic locations of DCM sites that we inferred as methylated (top panels in each pair) or unmethylated (bottom panels in each pair). The change in the DCM CCwGG methylation status is apparent as a shift in the distribution of the red curves at the A / T position outlined with the blue box. In black are reads from the control (unmethylated whole genome amplified DNA); in red are the native DNA signals. In many cases, the shift in signal is subtle. However, the identification of these sites as methylated or unmethylated is a binary classification of a continuous state - sites that we identify as unmethylated may in fact be methylated in 40% of all cells; sites we identify as methylated may be methylated in only 60% of all cells.

**Supplementary Figure S3.**
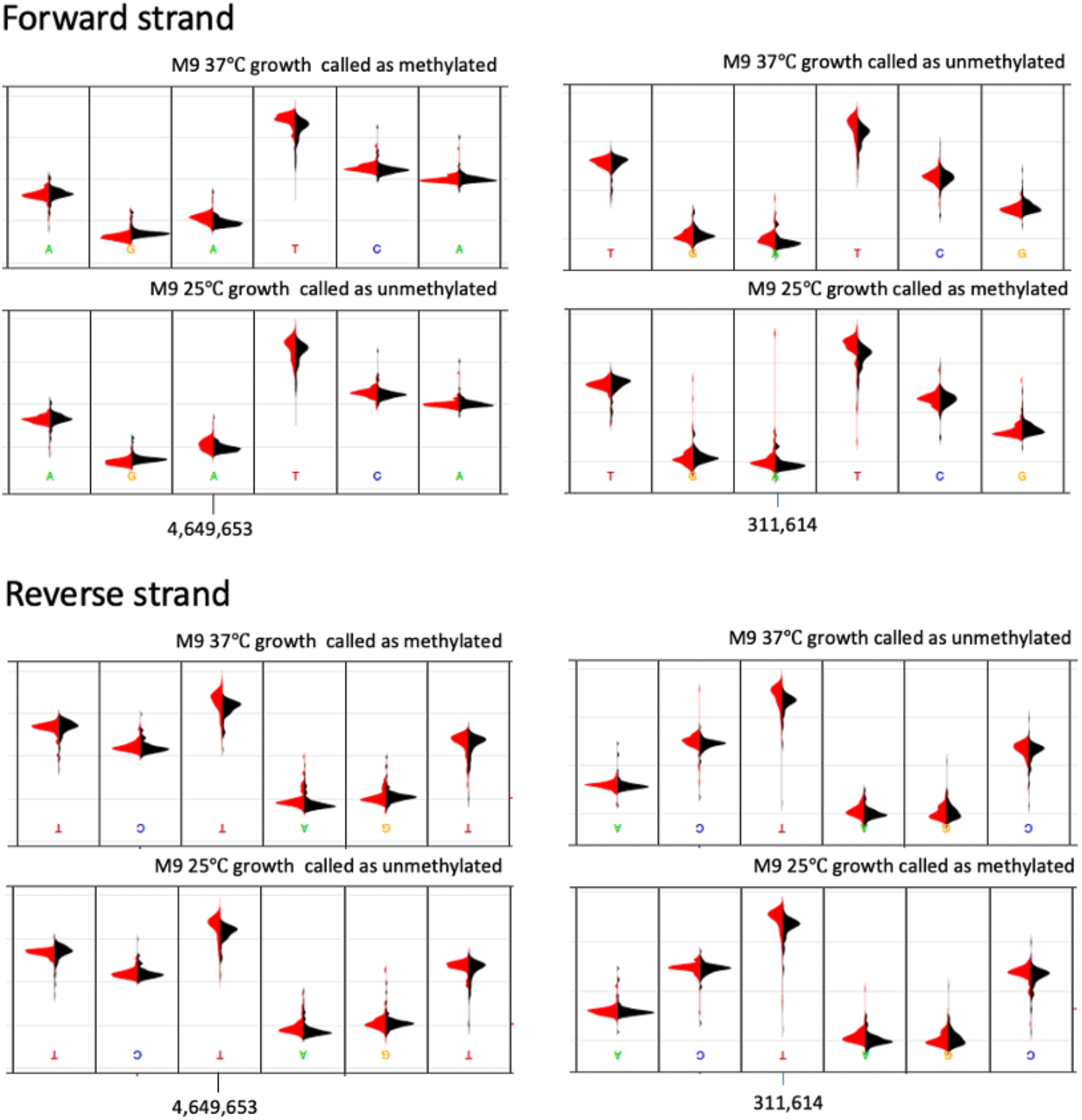
Identical DAM sites are inferred as methylated or unmethylated across different growth conditions. The change in the DAM GATC methylation status is apparent as a shift in the distribution of the raw nanopore signal from native DNA (red curves) at the T and A positions (the A is the modified base) compared to WGA unmodified DNA (black curves). Left panels: a DAM 6mA site that we inferred as methylated in M9 37ºC growth (top) but not during 25ºC growth (bottom). This is most apparent as a shift in the signal at the T position, for which the overlap between red and black is less in the top panel. Right panels: a DAM GATC site that we inferred as unmethylated in M9 37ºC growth (top) but methylated during 25ºC growth. Again, this is most apparent as a shift in the signal at the T position, with the overlap being higher in the top panel. Note that all native DNA molecules are not necessarily methylated at positions that we call as methylated, and vice versa: at positions that we call as unmethylated, all molecules are not necessarily unmethylated.

**Supplementary Figure S4.**
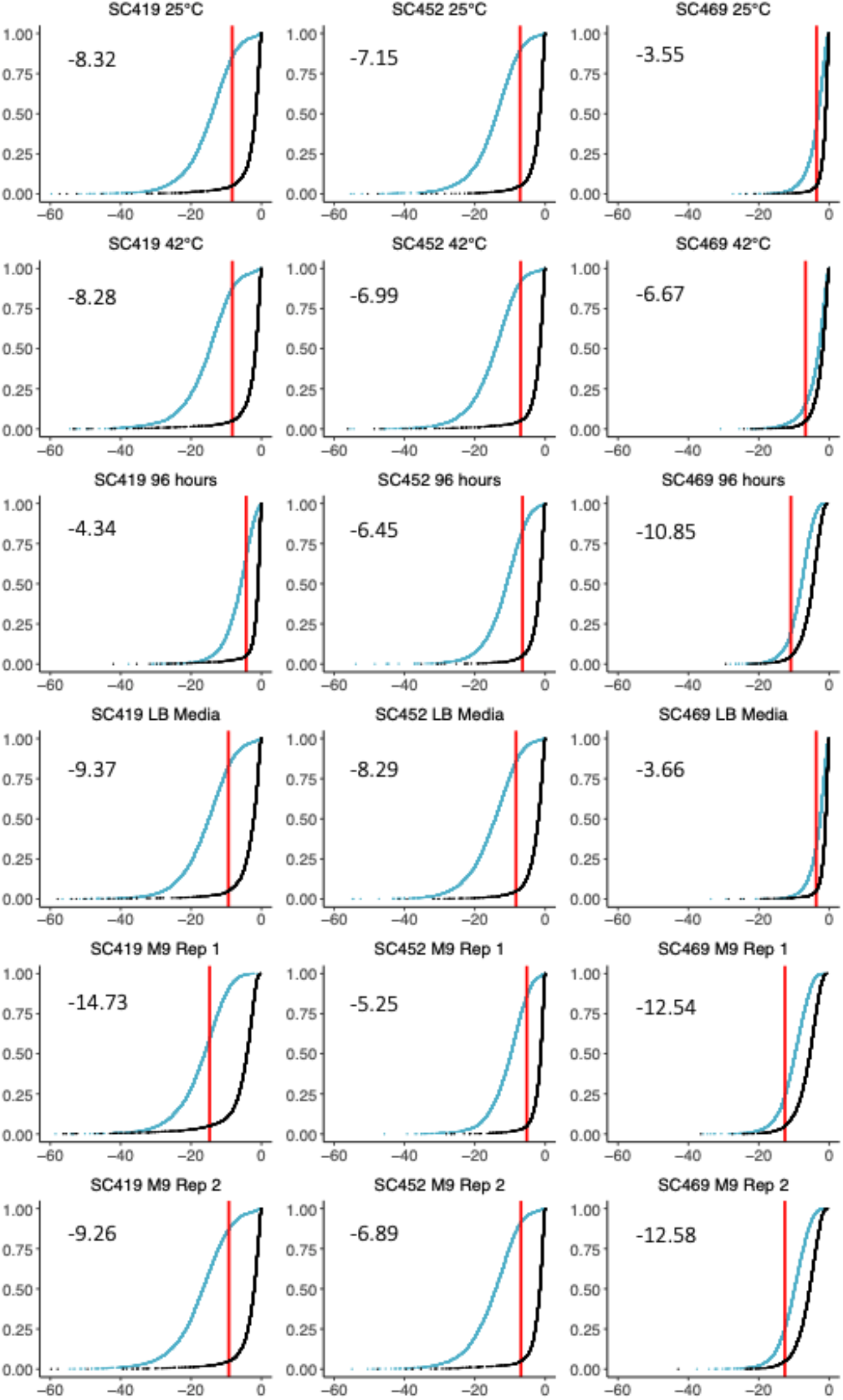
Cumulative distributions of p-values for DAM sites relative to random (unmethylated) sites. For each combination of isolate and growth condition we used the distribution of p-values at DAM binding sites (blue) and an equal number of random sites (black) to determine a p-value cut-off. This cut-off was established such that 10% of all unmodified sites were inferred as being modified, equivalent to a 0.1 FDR. Each cut-off is shown in red, and the log10 of the p-value cut-off is noted within each plot.

**Supplementary Figure S5.**
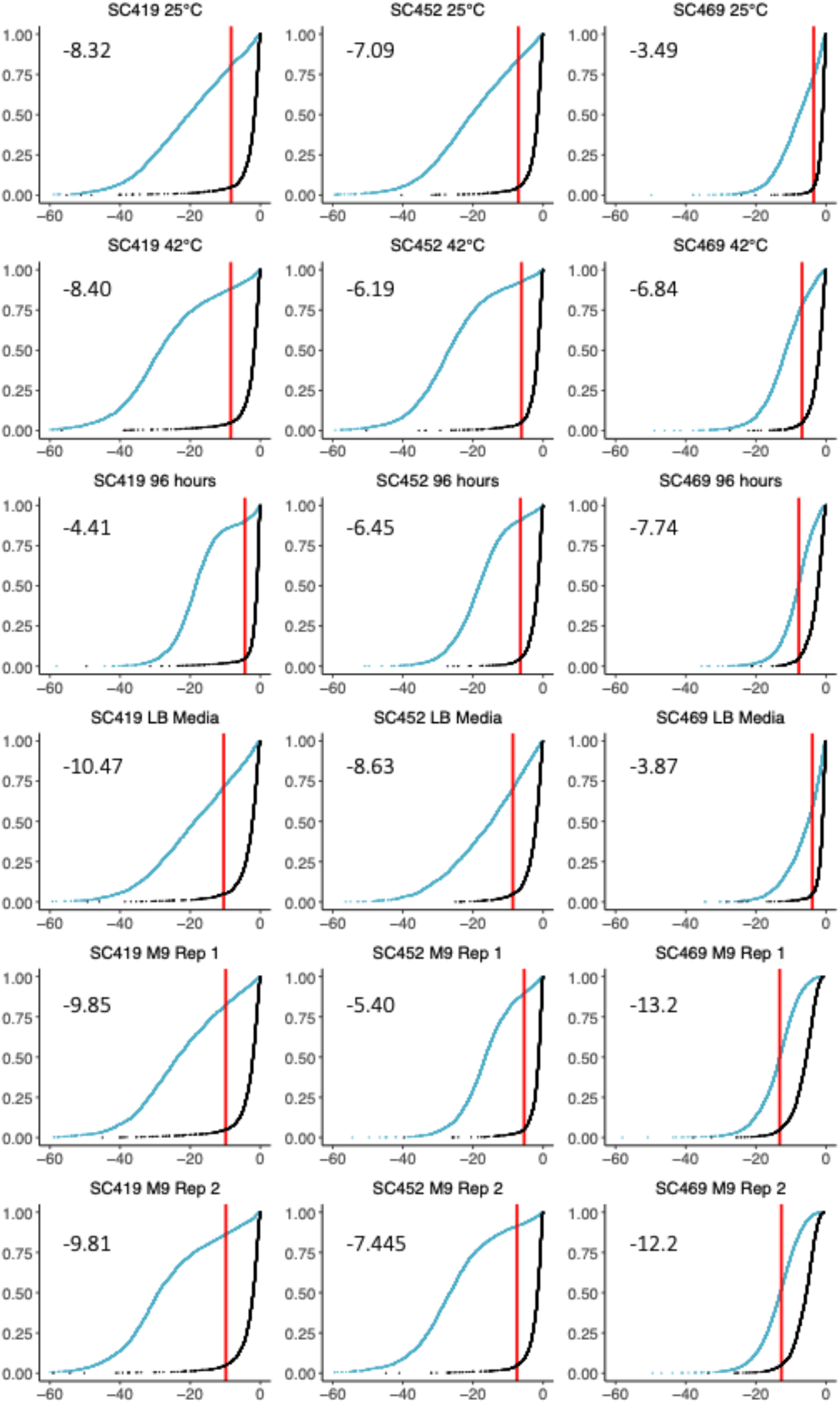
Cumulative distribution of p-values for DCM sites relative to random sites. For each combination of isolate and growth condition we used the cumulative distribution of p-values at DCM binding sites (blue) and an equal number of random sites (black) to determine a p-value cut-off equivalent to an FDR of 0.1. Each cut-off is shown in red, and the log 10 of the p-value cut-off is noted within each plot.

**Supplementary Figure S6.**
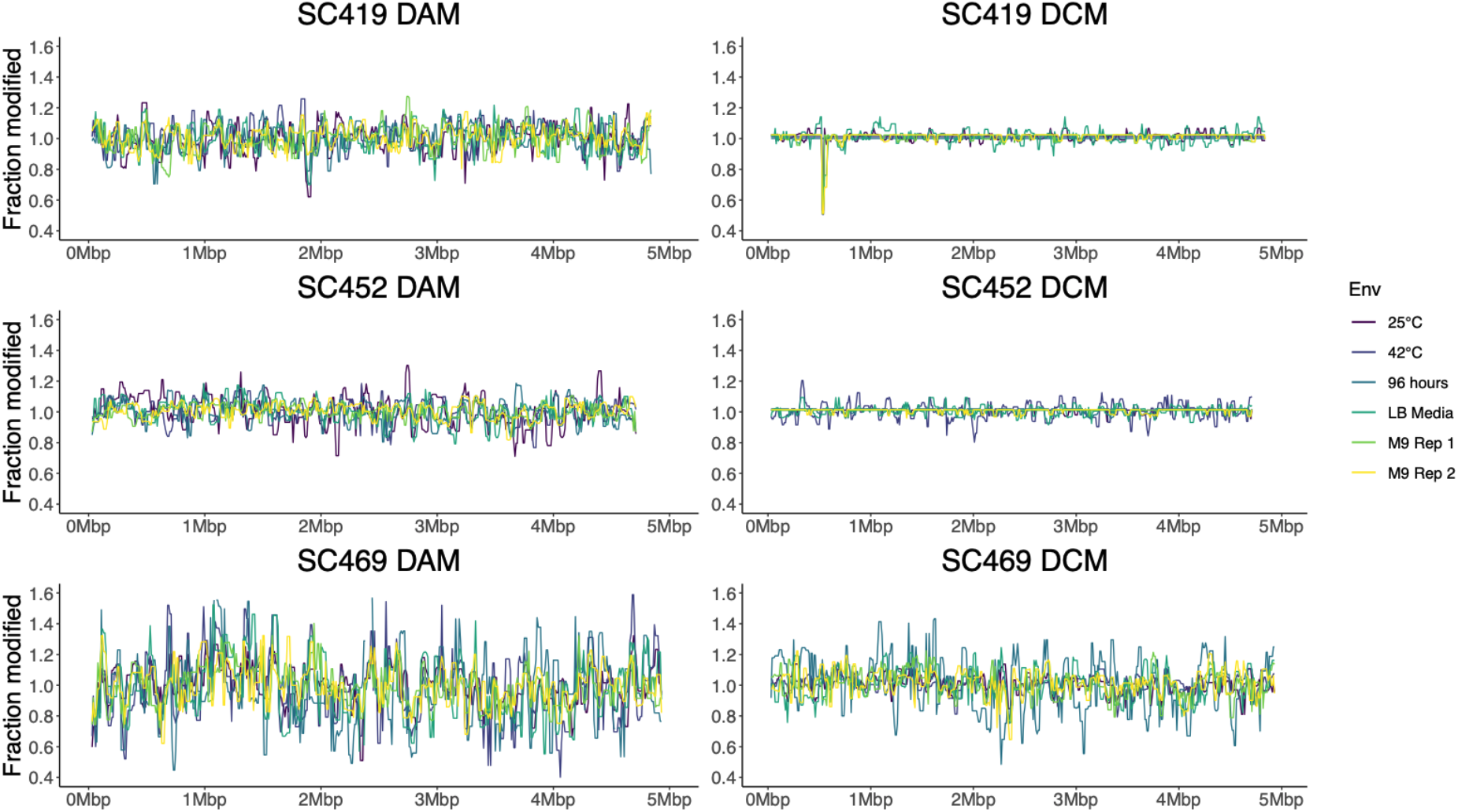
Mean-normalised fractions of modified sites across the genome. For each growth condition, we divided the fraction of modified sites in each window by the mean fraction of modified sites across all windows for that growth condition. This normalised fraction of modified sites are generally consistent across the genome for each methyltransferase and strain, which is clearly apparent in **Fig. 4**.

**Supplementary Figure S7.**
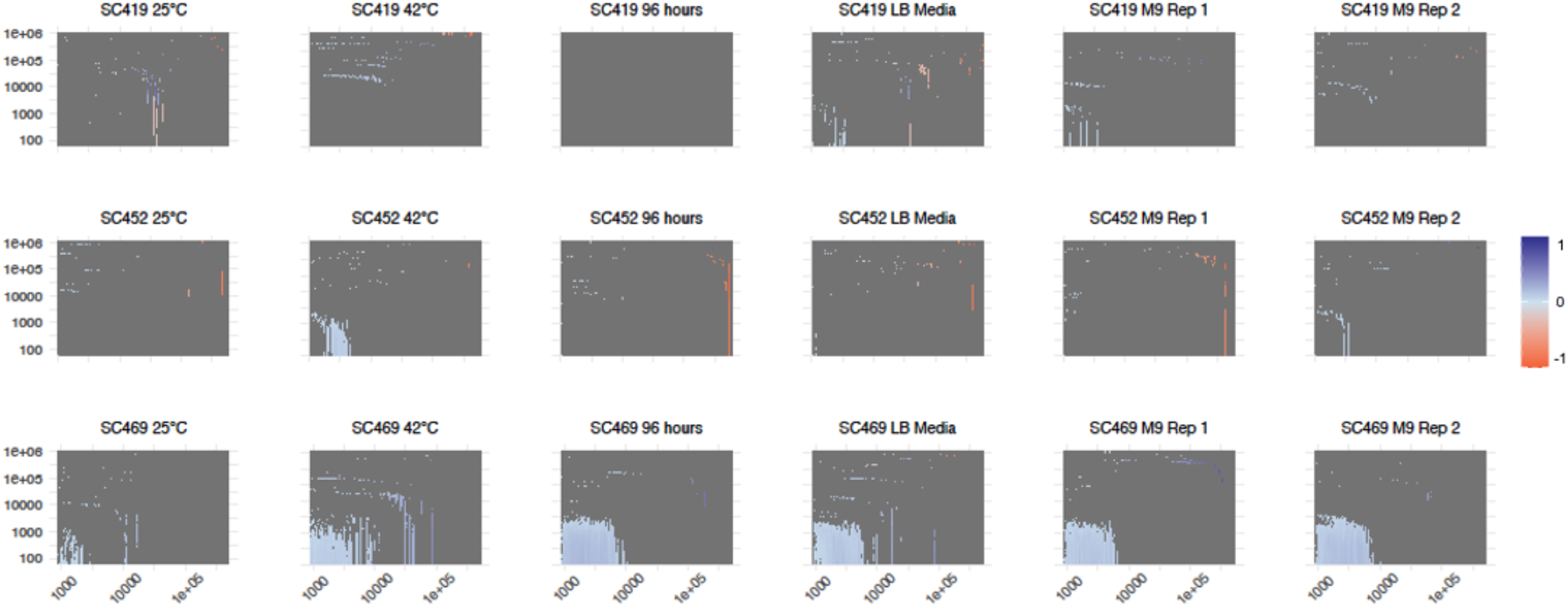
Global autocorrelation plots for DCM methylation. Each panel is a heatmap showing the correlation for the fraction of methylated DCM sites between windows of increasing size, ranging from 250 bp to 500 Kbp (different window sizes are plotted in columns), separated by increasing distances ranging from 0 (i.e., adjacent windows) to 1 Mbp (different distances are plotted in rows). Window sizes increase by a constant fraction of 4.7%; separating distances increase by a constant fraction of 9.6%. For example, the bottom left square in each heatmap shows the correlation in the fraction of methylated sites for neighbouring 250 bp windows; the middle square in each plot shows the correlation between 20.9 Kbp windows separated by 7.6 Kbp; the top right indicates 500 Kbp windows separated by 1 Mbp. In the example here, a standard autocorrelation function (ACF) would plot the correlations between windows of a certain size separated by a specific number of windows (e.g., 10 Kbp windows separated by 0 bp (neighbouring), 10 Kbp (one window), 20 Kbp (two windows), etc. This would be similar to several squares in the 53rd column in this plot: the squares in rows 1 (0 bp distance between windows), 56 (10 Kbp distance), 64 (20 Kbp distance), 68 (30.2 Kbp distance), 71, 74, and 76. However, this plot shows the analogous set of correlations at almost all window sizes and distances. For clarity only correlations with p < 0.01 are shown. In almost all cases, the correlations are positive (i.e., windows that are close tend to have similar levels of methylation), but this correlation only exists for windows up to approximately 5-8 Kbp in size and separated by a maximum of 5 Kbp. This suggests that there are no long-range correlations in the fraction of methylated sites. Note that the strongest correlations are observed for strain SC469, which is also the strain that exhibited the greatest variance in fraction methylated across genomic windows (**Fig. 3**). For other strains, the low level of variance in methylated fractions necessarily weakens the correlations.

**Supplementary Figure S8.**
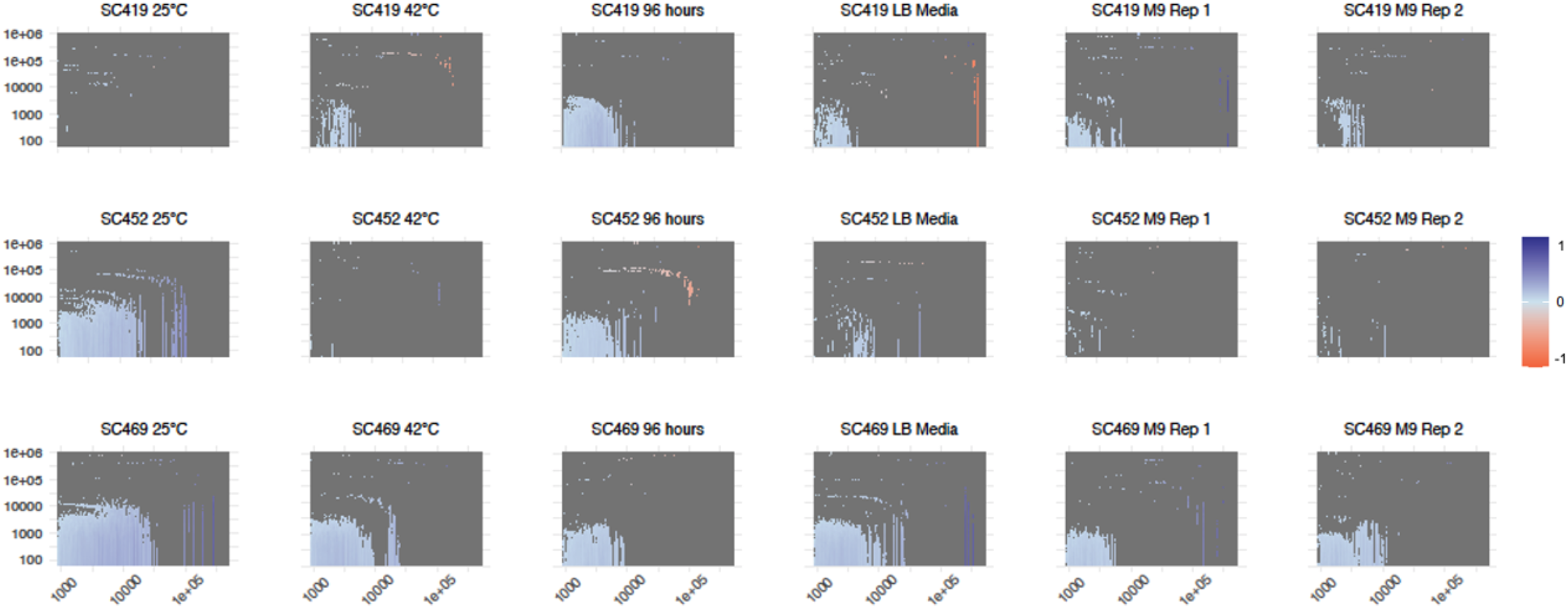
Global autocorrelation plots for DAM methylation. The annotation and details of this plot are the same as those shown in **Supp. Fig. S7** but for DAM methylation. Again, for clarity only correlations in p < 0.01 are shown. The correlations here in the fraction of methylated sites in a window are in general stronger but extend to a similar distance to those observed for DCM. Again, the strongest correlations are observed for strain SC469. However, correlations are also apparent for other strains in other conditions, also most likely because DAM methylated fractions exhibited much greater variation than DCM (**Fig. 3**).

